# Identification of new markers of angiogenic sprouting using transcriptomics: New role for RND3

**DOI:** 10.1101/2023.10.18.563021

**Authors:** Colette A. Abbey, Camille L. Duran, Zhishi Chen, Yanping Chen, Sukanya Roy, Ashley Coffell, Sanjukta Chakraborty, Gregg B. Wells, Jiang Chang, Kayla J. Bayless

## Abstract

**Background:** New blood vessel formation requires endothelial cells to transition from a quiescent to an invasive phenotype. Transcriptional changes are vital for this switch, but a comprehensive genome-wide approach focused exclusively on endothelial cell sprout initiation has not been reported.

**Approach and Results:** Using a model of human endothelial cell sprout initiation, we developed a protocol to physically separate cells that initiate the process of new blood vessel formation (invading cells) from non-invading cells. We used this model to perform multiple transcriptomics analyses from multiple donors to monitor endothelial gene expression changes. Single-cell Population Analyses, single-cell Cluster Analyses, and bulk RNA sequencing were used to delineate transcriptomic changes in invading cells. The results revealed a 39 gene signature that was consistent with activation of signal transduction, morphogenesis, and immune responses. Many of the genes were previously shown to regulate angiogenesis, and include multiple tip cell markers. Upregulation of SNAI1, PTGS2, and JUNB protein expression was confirmed in invading cells, and silencing JunB and SNAI1 significantly reduced invasion responses. Separate studies investigated Rounding 3 (RND3), also known as RhoE, which has not yet been implicated in angiogenesis. Silencing RND3 reduced endothelial invasion distance as well as filopodia length, fitting with a pathfinding role for RND3 via regulation of filopodial extensions. Analysis of *in vivo* retinal angiogenesis in *Rnd3* heterozygous mice confirmed a decrease in filopodial length compared to wild type littermates.

**Conclusion:** Validation of multiple genes, including RND3, revealed a functional role for this gene signature early in the angiogenic process. This study expands the list of genes that are associated with the acquisition of a tip cell phenotype during endothelial cell sprout initiation.

**HIGHLIGHTS:**

- Transcriptomic analyses identified 39 candidate genes that were upregulated at the onset of endothelial sprouting
- The gene signature includes signal transduction, morphogenesis, and immune responses
- Newly-identified RND3 is associated with filopodial extension and pathfinding

**Graphical Abstract:** 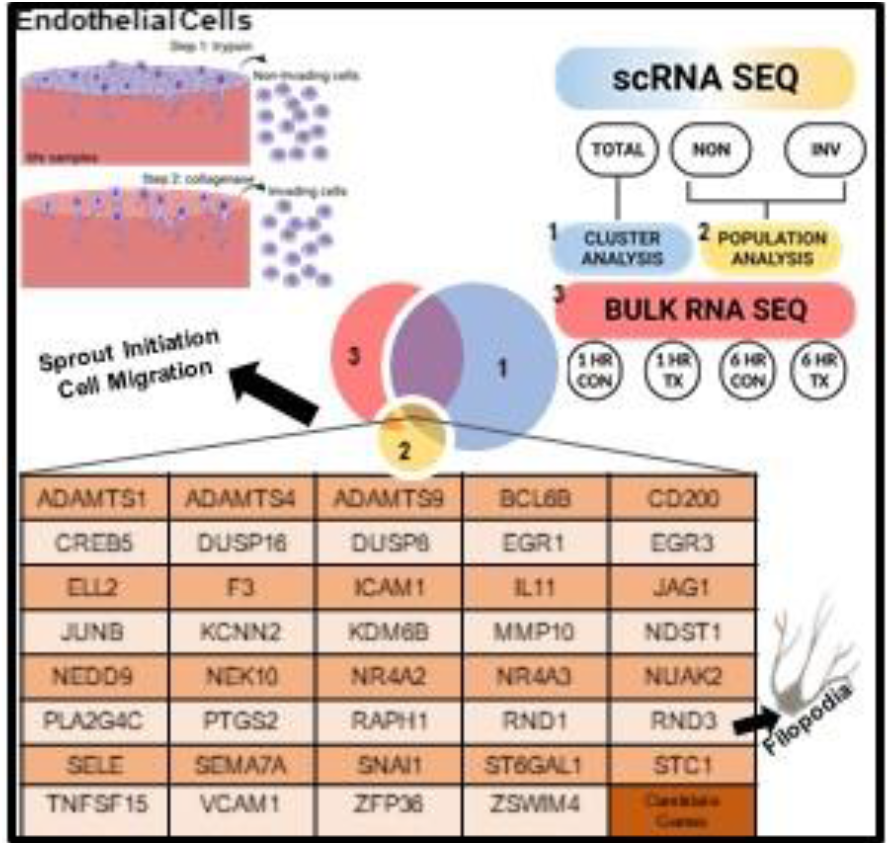

## INTRODUCTION

Angiogenesis is a multistep process involving careful coordination of distinct endothelial cell behaviors that ultimately accomplishes new blood vessel growth. Quiescent endothelial cells lining mature blood vessels are arranged as a single layer with stable junctions to maintain vascular integrity.^1^ In contrast, activated sprouting endothelial cells have vastly different characteristics.^2^ Given the known complexities in cell phenotypes,^3–5^ phenotype switching,^6–9^ and the abundance of signaling pathways involved in angiogenesis,^10–14^ gaining further insights into phenotypic changes displayed by individual endothelial cells during early sprout initiation will be invaluable to ultimately design effective strategies that antagonize, enhance, or normalize angiogenic vasculature.^12^

To a large extent, gene expression determines the identity, fate, functional response, and ultimately phenotype of individual cells.^15,16^ Unprecedented discoveries monitoring gene expression changes have been made possible by the development of high throughput sequencing (seq) technologies, including bulk RNA-seq and single-cell RNA-seq (scRNA-seq). Bulk RNA-seq analyzes large amounts of RNA from a single sample to discover differences in expression, splicing, sequence variation, and methylation events. One drawback, however, is that bulk RNA-seq does not distinguish between individual cells or cell types in a heterogeneous sample. This limitation has been overcome by scRNA-seq, which detects RNA transcripts from individual cells,^17,18^ even when originating from heterogeneous starting material.^19^ Both bulk and single-cell RNA-seq have provided insights into vascular development, heterogeneity and differentiation.^20–24^ In addition, endothelial cell expression profiles in lymph nodes^25,26^ and tumors,^27,28^ as well as during arteriovenous specification,^29^ coronary artery development,^30^ and various conditions of disease progression^4,31^ have been reported.

Transcriptional profiling provides an unprecedented opportunity to gain insights into genetic signatures that regulate biological processes. Our understanding of angiogenic sprouting has been aided by proteomics,^32–34^ knock-out and transgenic model organisms,^35–41^ and to a lesser degree transcriptomic analyses.^4,28,31^ To our knowledge, a comprehensive high resolution transcriptional analysis focusing solely on endothelial cell sprout initiation has not been reported. The current study applies both single-cell and bulk transcriptomic analyses for increased resolution of transcriptional changes that distinguish invading from non-invading endothelial cells. We discovered 39 candidate genes were commonly upregulated in activated, invading endothelial cells when comparing the transcriptomic data sets. The majority of these candidate genes have been previously implicated in angiogenesis, giving us confidence in the data set. Additionally, GO pathways most highly represented include signal transduction, morphogenesis, and immune evasion. Validation studies showed fundamental roles for multiple genes, including newly identified RND3. RND3 knockdown and haploinsuffiency in mice resulted in reduced filopodia length during retinal angiogenesis compared to WT littermates. Altogether, this study defines early events in endothelial sprout initiation more clearly by expanding the list of potential genetic markers upregulated early in the process of endothelial sprouting.

## METHODS

### Endothelial Cell Culture

Human umbilical vein endothelial cells from six independent donors (CC-2517/0000242397, C2517A/0000088928, C2517A/0000439577, C2517A/0000306191, CC-2517/0000198228, C2517A/0000699238: Lonza, TX) were used for transcriptomic analyses and validation. Cells were cultured at passages 3–6 in 75 cm^2^ flasks (Corning, NY) coated with 1 mg/mL sterile gelatin (Sigma,MO). Growth medium was previously described in detail ^42^ and consisted of M199 (Gibco, MA), supplemented with fetal bovine serum (Gibco, MA), bovine hypothalamic extract (Pel-Freeze Biologicals, AK), heparin (Sigma, MO), antibiotics (Gibco, MA), and gentamycin (Gibco, MA).

### scRNA-seq

Confluent human umbilical vein endothelial cells (HUVEC, passage 3-6) were used in invasion assays as previously described.^42^ Briefly, type 1 collagen isolated from rat tails (Pel-Freeze Biologicals, AK) was used to prepare collagen matrices at 2.5 mg/ml with 1 µM S1P (sphingosine 1-phosphate) (Sigma, MO) in 96 well half area plates (Costar, NY). The mixture (25 µl) equilibrated for 45 min, before seeding HUVEC (30,000 cells per well) in media containing M199 (Gibco, MA) supplemented with 1X RSII, ascorbic acid and 40 ng/ml bFGF and VEGF (R&D Systems, MN). For single-cell RNA-seq, cells were placed in an invasion assay for 6 hrs before non-invading, invading, and total cell populations were collected. *Non-invading cell collection*: Medium was removed, collagen matrices were rinsed with 100 µl of warm 1X Hepes Saline + 5 mM EDTA for 1 min at RT, 50 µl 0.25% trypsin-EDTA (Gibco, MA) was added per well, and cells were incubated at 37⁰C for 2-3 min. Plates were tapped to dislodge cells and 50 µl FBS (Invitrogen, MA) was added to each well to neutralize the trypsin. Liberated cells were collected, and wells were rinsed with 100 µl 1X Hepes Saline + 5 mM EDTA to collect the remaining non-adherent cells. Cells were centrifuged at 500 x g for 5 min and resuspended in 0.1% BSA in 1X PBS. *Invading cell collection*: Following removal of the non-invading cell population, the remaining collagen matrices (containing invading cells) were digested in M199 containing 1 mg/ml collagenase (Sigma, MO) at 37⁰C for 5 min. Digests were centrifuged at 500 x g for 5 min and resuspended in 0.1% BSA in 1X PBS. *Total cell collection*: At 6 hrs of incubation, the medium was removed, and matrices containing invading and non-invading populations were collected and digested in M199 containing 1 mg/ml collagenase at 37⁰C for 5 min. Samples were centrifuged at 500 x g for 5 min and resuspended in 0.1% BSA in 1X PBS.

### Collagenase Digestion vs. Direct Lysis

Endothelial cells seeded onto collagen matrices were collected after 6 hr of invasion into RLT lysis buffer directly (no collagenase exposure). Alternatively, collagen matrices were digested with collagenase, centrifuged, and then resuspended in RLT lysis buffer. The maximum time cells were exposed to collagenase was 10 minutes at 37°C.

### Generation of Single-cell Libraries

Single-cell libraries for each sample were generated at the Texas A&M Institute for Genome Sciences and Society (College Station, Texas). Single-cell suspensions collected using the above protocol were loaded in the 10X Chromium Controller using the Chromium Single-cell 3’ v3 Kit per the manufacturer’s protocol. All samples were processed in the same batch. Sequencing was performed with Illumina NextSeq 500.

### Computational Analysis of Single-cell Libraries

The scRNA-seq analyses were performed with the 10X Genomics Cell Ranger (v4.0) pipeline using the default settings. Cell Ranger computations were performed by the Texas A&M Institute for Genome Sciences and Society core facility. The raw Illumina NextSeq 500 base call file outputs were demultiplexed into FASTQ files by use of *cellranger mkfastq*. Then, *cellranger count* was used to align reads to the human genome GRCh38 and generate single-cell feature counts.

For the Population Analysis, libraries from two different cell donors of non-invading and invading populations were aggregated and read depth was normalized using *cellranger aggr*. The same method was applied for the two cell donors aggregated in the total sample Cluster Analysis. Loupe Cell Browser (v6.0.) was used to filter cells with high mitochondrial gene expression (10 or 15 percent threshold) and excessively high or low UMI counts. For the aggregated biological replicates, Loupe was used to calculate differential gene expression, producing relative comparisons of gene expression between non-invading and invading populations. To regress the impact of collagenase exposure, cells with FOS expression > 0 were excluded using Loupe prior to clustering. Cluster Analysis was done on the total samples using Kmeans clustering. The clusters representing the invading and non-invading cell populations, as defined by the Cluster Analysis, were compared. Per the default methods of Loupe, p-values were calculated using the sSeq variant of the negative binomial exact test and were corrected for false discovery rate (FDR) using Benjamini-Hochberg method for multiple comparisons.^43^ An FDR-corrected p-value threshold of 0.1 was used to determine statistical significance in the single-cell analysis.

### Bulk RNA-seq

Samples analyzed for bulk RNA sequencing included control and activated cells. Control (non-activated) cells were collected after seeding onto collagen matrices without S1P, bFGF or VEGF for 1 and 6 hrs (1 HR CON, 6 HR CON). Invading (activated) cells were treated with S1P and growth factors identically to scRNA-seq experiments prior to collection at 1 and 6 hours (1 HR TX, 6HR TX). We have previously shown that endothelial cells activated with both S1P and growth factors undergo robust invasion.^44–46^ Thus, the control (CON) groups at 1 and 6 HR consist of non-invading cells, while invading cells are present in the treated (TX) group only.^44,45^ HUVEC from three individual donors were used for these experiments. At the specified time point, media was removed, and samples were digested in 1 mg/ml collagenase/M199 and incubated at 37⁰C for 5 min. Cells were centrifuged at 500 x g for 5 min. Supernatant was removed, and cell pellets were resuspended in RLT lysis buffer.

RNA was extracted, treated with DNase on the column using Qiagen’s RNeasy Kit per manufacturer’s instructions, and processed using Texas A&M AgriLife Research Genomics & Bioinformatics Services for sequencing using a 125-PE HiSeq platform with Illumina’s TruSeq RNA Library Prep. STAR2.3.1 was used to align the reads to the reference genome hg19 with mapping rates between 93%-94%. Differential expression analysis using DESeq was used to perform two comparisons: 1 HR CON vs. 1 HR TX, and 6 HR CON vs. 6 HR TX to compare mRNA expression between CON (non-invading) and TX (invading) populations.

### Validation of mRNA Expression Using qPCR

Invading and non-invading cells were collected after 6 hrs of incubation using the same method as described for scRNA-seq. Cell pellets were resuspended in RLT Lysis Buffer, and RNA was extracted with Qiagen’s RNeasy Kit. Additionally, total gels were collected at 1, 3 or 6 hrs of invasion for qPCR analysis of gene expression during siRNA invasion assays to confirm gene knockdown and evaluate expression profiles. cDNA was made using Bio-Rad’s iSCRIPT cDNA synthesis kit with 0.5-1 µg RNA as starting template. Real-time PCR was performed using iTaq Universal SYBR Green Supermix (Bio-Rad, CA), 0.5 µM forward and reverse primers and 1 µl cDNA template, diluted 1 to 100, on a StepOnePlus Real-time System (ABI, MA) for data analysis as 2-^ΔΔCT^. RPLP0 was used as the housekeeping gene. Primer sequences are listed in Supplemental Table S1.

### Immunofluorescence and Imaging

Invading cultures at the time points indicated were fixed in 4% paraformaldehyde (Electron Microscopy Sciences, PA) for 20 min, washed twice in 25 mM Tris with 200 mM glycine for 15 min, permeabilized with 0.5% Triton X-100 in PBS for 30 min, and incubated in blocking buffer (0.1% Triton X-100, 1% BSA, 0.2% sodium azide, 5% goat serum) at 4°C overnight. Primary antibodies were used at 1:200 in blocking buffer for 3 hr at RT and samples were washed 4 times for 30 min each in 0.1% Triton X-100 in PBS. Goat anti-rabbit or anti-mouse secondary antibodies conjugated to Alexa 488 or Alexa 594 (Invitrogen, MA) were added (1:600 dilution in blocking buffer) for 1 hr at RT. Samples were washed twice for 30 min in 0.1% Triton X-100 in PBS and again overnight in 0.1% Triton X-100 in PBS. The next morning, a final wash containing 10 µM DAPI was performed. Confocal images were captured with a Nikon Eclipse TI equipped with NIS Elements AR software. Antibodies are listed in the Major Resources Table. For quantifying JUNB nuclear intensity, image stacks were collected to identify invading and non-invading cells. JunB nuclear intensity was determined by identifying the center of the nucleus and recording JunB and DAPI signaling intensities. Data were separated into invading and non-invading groups for analysis.

### siRNA Transfection

Silencer select siRNAs (Ambion 4392420) to SNAI1: ID# s13187: CACUGGUAUUUAUAUUUCAtt; β2M: ID# s1852: GAAUGGAGAGAGAAUUGAAtt; RND3: ID# s1577: GAACGUGAAAUGCAAGAUAtt, ID#s1579: GAUCCUAAUCAGAACGUGAtt and Negative Control #2 (Cat# 4390846) were used at 50-100 nM to transfect HUVEC (P3-P6) that were seeded at 50% confluency onto gelatin (1 mg/ml) coated flasks with siPORT Amine (Ambion, MA) per manufacturer’s instructions. Cells were used in invasion assays and collected for Western blotting and RNA extraction 48-72 hrs after transfection.

### ASO Transfection

Custom ASOs were designed for JUNB (5’-TCTGGCGCGATAGCTT-3’) and a negative control scramble (5’-AACACGTCTATACGCG-3’). HUVEC, passage 3, were transfected using siPORT Amine (Ambion, MA) with 50 nM ASO, per manufacturer’s instructions. Briefly, P3 HUVEC were seeded onto gelatin-coated flasks (1mg/ml) at 90% confluency. The next day, cells were transfected with 12.5 µl siPORT Amine and 50 nM ASO in 3 ml OMEM (no antibiotic) containing 5% FBS. After overnight incubation, cells were rinsed with OMEM (no antibiotic) and fed with antibiotic free growth media. The following day, cells were used in invasion assays and extracts were collected for Western blotting and for RNA extraction.

### Evaluation and Quantification of Invasion Responses

Overnight invasion cultures were fixed in 3% glutaraldehyde (Sigma, MO) and stained with 0.1% toluidine blue in 30% methanol. Toluidine blue-stained collagen matrices were cut and imaged from the side at 4X on an Olympus CKX41 microscope fitted with a Q Color 3 camera to document invasion responses. Separate samples were stained with DAPI prior to capturing confocal images with a Nikon Eclipse T1 microscope and rendered in the volume view with Alpha depth coding selected. Nuclear invasion distance was quantified using NIS Elements AR software. A minimum of 25 structures from three separate experiments were analyzed. Data shown are from a representative experiment.

### Protein Extracts and Western Blotting

To quantify protein levels in cells, pellets were lysed directly with hot 1.5X Laemmli sample buffer with 2% mercaptoethanol. Protein extracts were incubated at 95°C for 5 min before loading onto SDS-PAGE gels (7-12%) for electrophoresis. Protiens were transferred to 0.45 µm Protran Membrane (GE, IL) for 90 min at 140V. Membranes were blocked for 1 hr in 5% milk before addition of primary antibodies and incubated at 4°C overnight. Blots were washed 3 times (5 min per wash in Tween-20 saline) before addition of HRP-conjugated secondary antibodies (1:5000). After a 1 hr incubation with the secondary antibody, blots were again washed 3 times (5 min per wash in Tween-20 saline) before addition of ECL (Millipore, MA) for chemiluminescent detection on a BioRad Image Analyzer using Image Lab to capture and quantify signal intensity. Densitometry values were obtained using Image Lab software to normalize signal intensity of investigated proteins to GAPDH or CD31 as indicated. Analyses were performed with at least three independent replicates. Antibodies are listed in the Major Resources Table. Uncropped Western Blots are included in Supplemental Fig. S1.

### Retinal Angiogenesis Assays

Rnd3 haploinsufficient mice were generated as previously reported.^47^ Briefly, the targeting vector was inserted at Rnd3 intron 2 in the ES cell line with C57BL/6N background. The chimeras were inbred with wild-type C57BL/6J (Jackson Laboratory) for more than 30 generations. The age-matched wild-type littermates were used as control. Retinas were isolated at postnatal day 5 and stained with 10 μg/ml FITC-conjugated lectin from *Griffonia simplicifolia* (isolectin-B4)^48^ (L9381, Sigma, MO). Images were collected from each lobe and analyzed by an observer blinded to mouse genotype. Data from at least 5 fields per retina were pooled and averaged to provide a value for each retina.

### GOnet Pathway Analysis

GOnet software (http://tools.dice-database.org/GOnet/) was used for GO term annotation analysis on the list of commonly upregulated genes in invading endothelial cells by selecting bio processes with go annotation in csv output.

### Statistical Analysis

GraphPad Prism version 6.07 for Windows (GraphPad Software, La Jolla California USA, www.graphpad.com) was used to perform all statistical analyses. Raw data values were analyzed for normality and equal variation prior to application of parametric or non-parametric statistical tests. Individual tests applied are indicated in each figure legend, along with exact P-values. Error values in all data sets represent standard deviation.

## RESULTS

### Three-Dimensional Assays Allowed Separation of Invading and Non-Invading Endothelial Cell Populations for Single-cell RNA Sequencing

A three-dimensional model of primary human endothelial sprout initiation was employed to find genes important for initiation of angiogenesis. Cells were seeded as a monolayer onto a 3D collagen matrix and supplied with pro-angiogenic factors to induce sprouting. After 24 hours of incubation a population of endothelial cells have completed the invasion process (Fig 1A). We chose the 6hr time point as the cells have just begun the invasion process, and we hypothesized that the genes important for the initiation of sprouting angiogenesis/angiogenic switch would be upregulated at this time. At 6 hours, non-invading and invading cells are clearly distinguishable, physically separable, and the latter extend processes into collagen matrices at this time point (Fig. 1A). We performed both single-cell and bulk RNA transcriptomics to identify changes in gene expression between non-invading and invading cells. In scRNA-seq experiments, two independent donors were used to collect separate samples of non-invading (NON) and invading (INV) cells for a Population Analysis (Fig. 1B). Additionally, samples containing all invading and non-invading cells (TOTAL) were collected and evaluated using scRNA-seq Cluster Analysis. The schematic in Fig. 1C illustrates the separation of non-invading and invading populations used for scRNA-seq. The non-invading population was liberated by treating the monolayer of 6 hr cultures with trypsin. With the non-invading population removed, invading cells were collected from collagen matrices by collagenase treatment. Photographs of cell monolayers (Supplemental Fig. 2A) and a side view of invading cells (Supplemental Fig. 2B) before and after trypsinization reveal that the invading cells that extended processes into collagen matrices remain behind. This method was used to collect a viable sample of single cells from non-invading (NON) and invading (INV) populations suitable for scRNA-seq (Supplemental Fig. 2C).

**Fig. 1.**
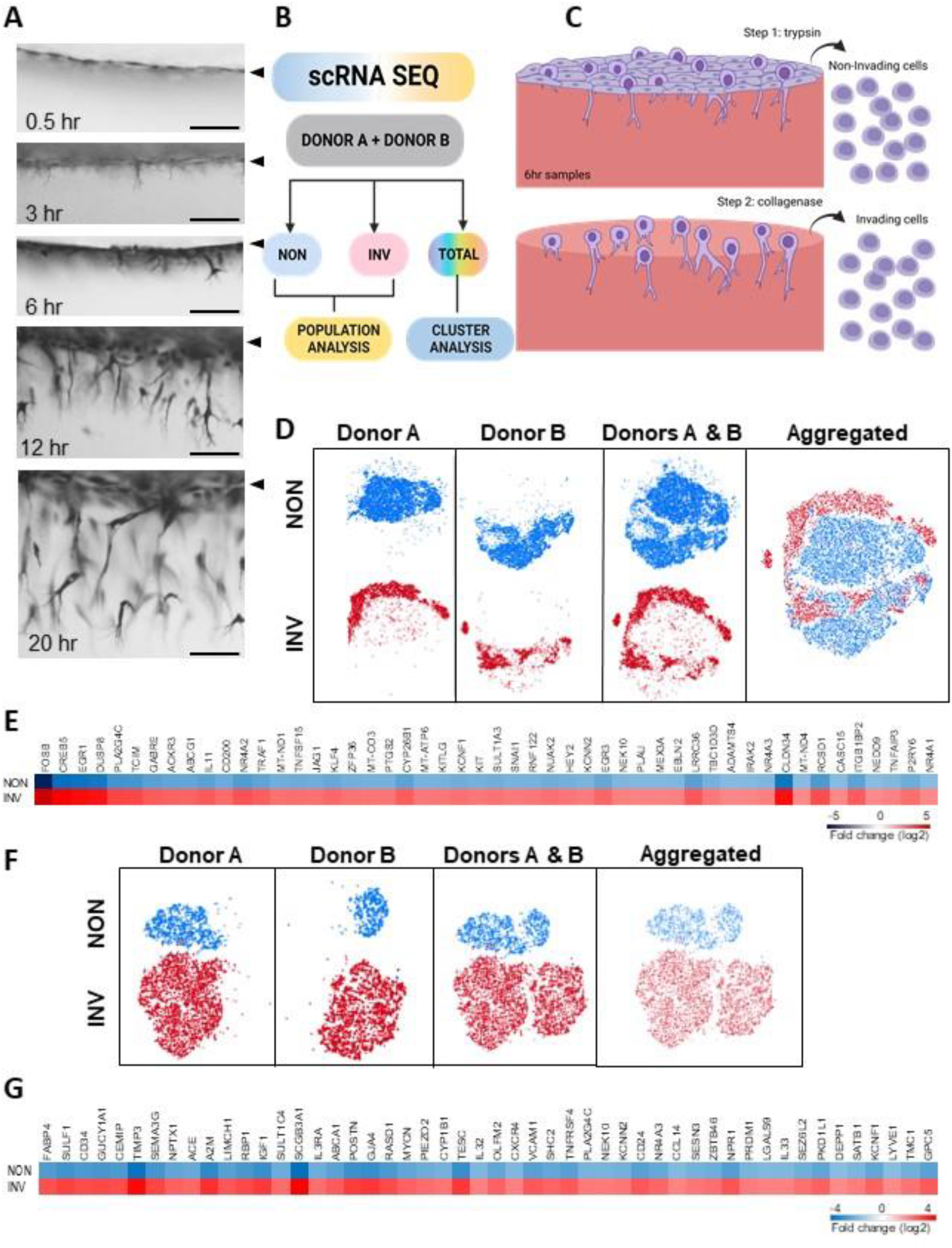
Overview of sample collection and methods to separate non-invading and invading endothelial populations from three-dimensional assays. **(A)** ECs were allowed to invade a 3D collagen matrix in the presence of S1P, VEGF and bFGF. Photographs of side views of invasion responses at the times indicated. Samples were fixed and stained with toluidine blue. Arrowheads indicate the original monolayer. Scale bars in all panels = 100 µm. **(B)** Flow chart depicting the multi-faceted approach used to determine genes upregulated in invading cells. For scRNA-seq, samples from two independent donors (A and B) were sequenced separately for non-invading (NON), invading (INV), and total cell populations. **(C)** Graphical representation of workflow to separate non-invading and invading populations for scRNA-seq analyses. Monolayers of intact cultures were treated with trypsin to liberate non-invading cells. Invading cells extending processes into collagen remained behind and were digested with collagenase. Artwork created with BioRender.com **(D)** t-SNE plots for Population Analysis of NON (blue) and INV (red) cells at 6 hr, with each lot displayed separately and aggregated. (E,G) Heatmaps displaying expression of top 50 markers upregulated in the INV clusters for Population and Cluster Analyses, respectively. **(F)** t-SNE plots for cluster analysis of NON (blue) and INV (red) cells at 6 hr, with each lot displayed separately and aggregated. t-SNEs plots show precise cluster groups of NON (blue), INV (red) and Aggregated data from both biological replicates.

### Collagenase Disassociation Artificially Induced FOS Expression

The initial scRNA-seq Population Analysis comparing non-invading to invading endothelial cells revealed substantial upregulation of various early response genes (ERGs) including *EGR1, FOS, FOSB, JUNB and ZFP36*. Recent reports have shown disassociation of tissue with collagenase can cause spurious gene expression of ERGs.^49–51^ Due to this potential artifact in our expression data, we compared direct lysis of invading cells versus 10-minute collagenase exposure before lysis. A qPCR analysis revealed that *FOS* expression was substantially upregulated with collagenase digestion compared to cells not exposed to collagenase but other ERGs evaluated were not (Supplemental Fig. 3A). Based on this result, FOS expressing cells were excluded from single-cell data sets and re-clustered. In doing so, 5394 and 4117 cells were removed from the aggregated Population and Cluster Analyses, respectively before proceeding to additional downstream analyses. A summary of scRNA-seq parameters metrics, including number of cells analyzed, and reads per cell, is shown in Supplemental Table S2.

### Population Analyses Derived From scRNA-Seq of Invading and Non-Invading Cell Populations Revealed Significant Upregulation of 180 Genes in the Invading Population

We first performed a Population Comparison of non-invading and invading endothelial cells which generated 180 hits (Supplemental Table S3). Precise clustering of the population of NON (blue), INV (red), and aggregated (red and blue), from both biological replicates (Donors A and B) revealed that single-cell sequencing yielded two populations with distinct transcriptional expression profiles (Fig. 1D). These data show successful separation of invading from non-invading endothelial cells for downstream scRNA-seq analysis. A heat map of the top 50 differentially expressed genes from the Population Analysis is shown (Fig. 1E).

### Cluster Analyses Derived from scRNA-Seq of a Total Cell Population Identified an Invading Cluster Enriched with Tip Cell Markers

A second transcriptomic analysis used total samples containing a mix of invading and non-invading cells (Fig. 1B). A Cluster Analysis was performed to determine if invading and non-invading populations could be efficiently identified from a mixed population using scRNA-seq. Guided by the gene set obtained from the Population Comparison that assigned markers to INV and NON groups (Supplemental Table S3), we identified a non-invading cluster (blue) and an invading cluster (red) from total samples (Fig. 1F). This analysis generated 1176 differentially expressed genes with upregulated expression in the invading cluster. Supplemental Table S4 shows a complete list of the local Cluster Analysis comparison between the INV (red) and NON (blue) clusters. A heat map of the top 50 differentially expressed genes from the Cluster Analysis is shown (Fig. 1G). We found the entire invading cluster is enriched with multiple tip cell markers, including *CD34*, *CXCR4*, *DLL4,* and others, giving us confidence in our experimental design and analysis to identify markers of angiogenic sprouting. We also observe that cells are not substantially segregating based on library of origin, but rather on expression profiles categorizing them as either invading or non-invading (Fig. 1F).

### Bulk RNA Sequencing of Invading and Non-Invading Cell Populations Revealed Early Transcriptional Changes Consistent with a Sprouting Phenotype

Bulk RNA sequencing (bulk RNA-seq) experiments using three independent endothelial cell donors were performed to complement the scRNA-seq analysis (Fig. 2A). Experimental groups included non-activated control samples (CON) and samples treated with pro-angiogenic stimuli that were activated (TX). The CON group showed no invasion, while the TX group showed robust invasion initiation responses after 1 and 6 hrs of incubation^46^. RNA was collected from non-activated and thus an entirely non-invading cell population (1 HR CON and 6 HR CON) for comparison with activated samples that contained a mixture of invading and non-invading cells (1 HR TX and 6 HR TX). Cells were collected and sequenced, and the relative expression of individual genes was determined. The summary of bulk RNA-seq metrics and genes identified from the bulk RNA-seq analysis are included in Supplemental Tables S5 and S6, respectively. This analysis generated 453 differentially expressed genes that were significantly upregulated at 1 or 6 hr of invasion in treated versus control samples. No adjustments for collagenase treatment were included, as both the control and treated cell populations were collagenase digested. A heat map of the top 50 differentially expressed genes from the bulk RNA-seq analysis is shown (Fig. 2B).

**Fig. 2.**
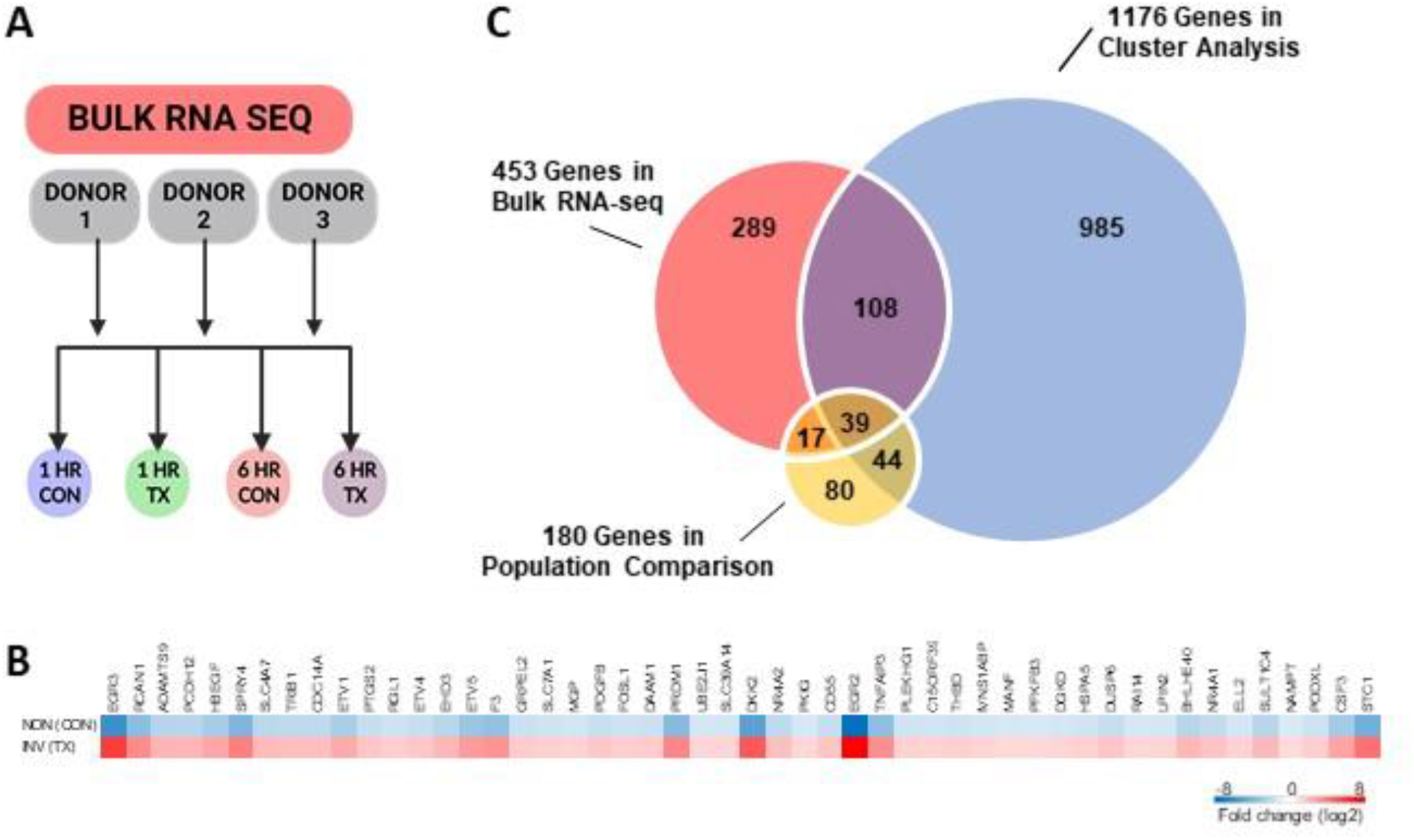
Integration of Population, Cluster, and Bulk RNA-seq Analyses revealed 39 candidate genes consistently upregulated in invading cells. **(A)** Bulk RNA-seq data was acquired from total samples of three independent donors (1-3) and evaluated for differential expression. Groups collected include non-activated and non-invading controls at 1 and 6 hr (1 HR CON, 6 HR CON) and activated, treated and invading cells at 1 and 6 hrs (1 HR TX, 6 HR TX). Artwork created with BioRender.com **(B)** Heatmap displaying expression of top 50 markers upregulated in the invading (INV) clusters for bulk-RNA sequencing. **(C)** Venn diagram depicting the number of upregulated genes in invading samples categorized by the analysis method indicated. Red: bulk RNA-seq; Blue: scRNA-seq cluster analysis; Yellow: scRNA-seq population comparison. Note 39 genes are identified at the intersection of the three analysis methods.

### Integration of Population, Cluster, and Bulk RNA-Seq Analyses Revealed 39 Candidate Genes Consistently Upregulated with Invasion

To establish a unique and validated list of genes upregulated in sprout initiation, we integrated information from the scRNA-seq Population Comparison (yellow), Cluster Analyses (blue) and bulk RNA-seq data (red) and looked for overlap among the data sets (Fig. 2C). The Venn diagram shows that 39 candidate genes were commonly upregulated in all three analyses (Fig. 2C). The Cluster Comparison yielded 83 genes in common with the Population Comparison. Considering the bulk RNA-seq comparisons, 56 of the 180 genes were upregulated in activated, invading cells at either 1 hr or 6 hrs. The 39 candidate genes ultimately common to all three data sets are listed in Table 1. A comparison of all data sets can be found in Supplemental Table S7. Of these, we observed genes that have been reported to play a role in angiogenic responses, as well as those uninvestigated in angiogenesis (Table 1). To ensure that collagenase digestion was not responsible for elevated expression of these candidate genes, we chose one gene from each category in Table 1 and performed qPCR on samples that were extracted with or without collagenase as in Figure 2. We did not see a significant increase in expression of the candidate genes selected in response to collagenase treatment (Supplemental Fig. 3B), indicating that changes in gene expression are associated with alterations in phenotype between invading and non-invading cells. Ultimately, these analyses converged to 39 commonly upregulated transcripts found in invading endothelial cells.

**Table 1.**
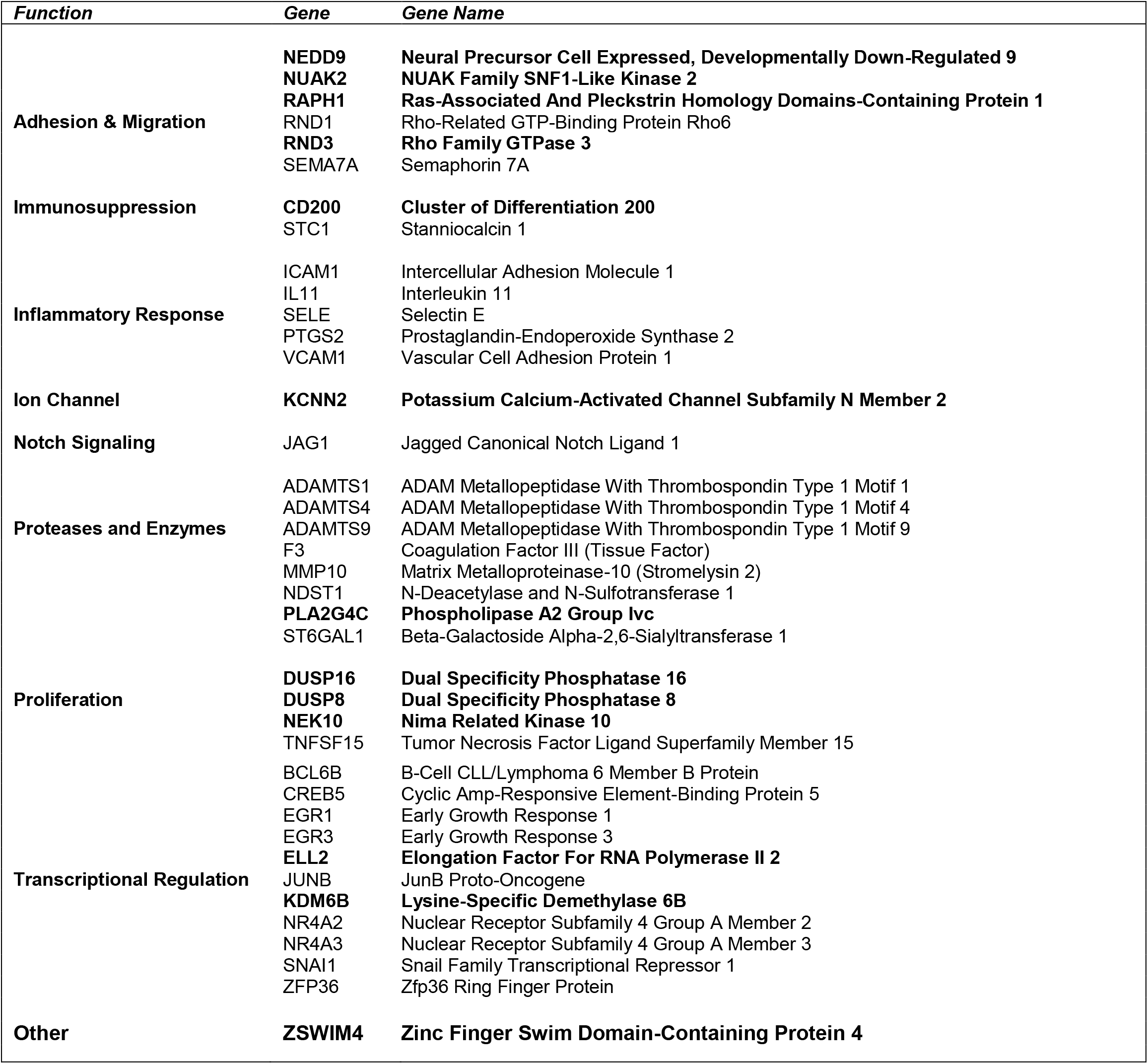
Candidate genes consistently upregulated in invading cells. Genes are listed alphabetically in functional categories based on GO terms, identified by gene symbol and name. Genes in bolded text have not been shown to play a role in angiogenic responses. See Supplemental Table S8 for supporting citations.

### GO Pathway Analysis Identified Expected Changes in Signal Transduction and Morphogenesis Along with a Strong Pro-Inflammatory Profile

Analysis of the genes commonly upregulated with endothelial invasion revealed the majority regulate signal transduction and developmental processes (Fig. 3A). Interestingly, almost half (46%) of the commonly upregulated genes are also involved in immune system processes, including *BCL6B*, *CD200*, *EGR1*, *EGR3*, *ICAM1*, *IL11*, *JAG1*, *JUNB*, *KDM6B*, *NEDD9*, *NR4A3*, *SELE*, *SEMA7A*, *ST6GAL1*, *STC1*, *TNFSF15*, *VCAM1*, and *ZFP36*. Differential gene expression of immune related transcripts between INV and NON cells in the Population, Cluster, and bulk RNA-seq Analyses is depicted with heat maps in Figure 3B. A stark upregulation in expression of the genes associated with immune regulation and inflammatory responses are displayed from the bulk RNA-seq data, where increased expression occurs with activation at either 1 and/or 6 hours of invasion (Fig. 3C). Together, these data reveal anticipated changes in signaling processes known to be associated with angiogenic responses, such as signal transduction, differentiation, and motility, as well as unexpected previously unrecognized upregulation of markers of inflammation and immune evasion.

**Figure 3.**
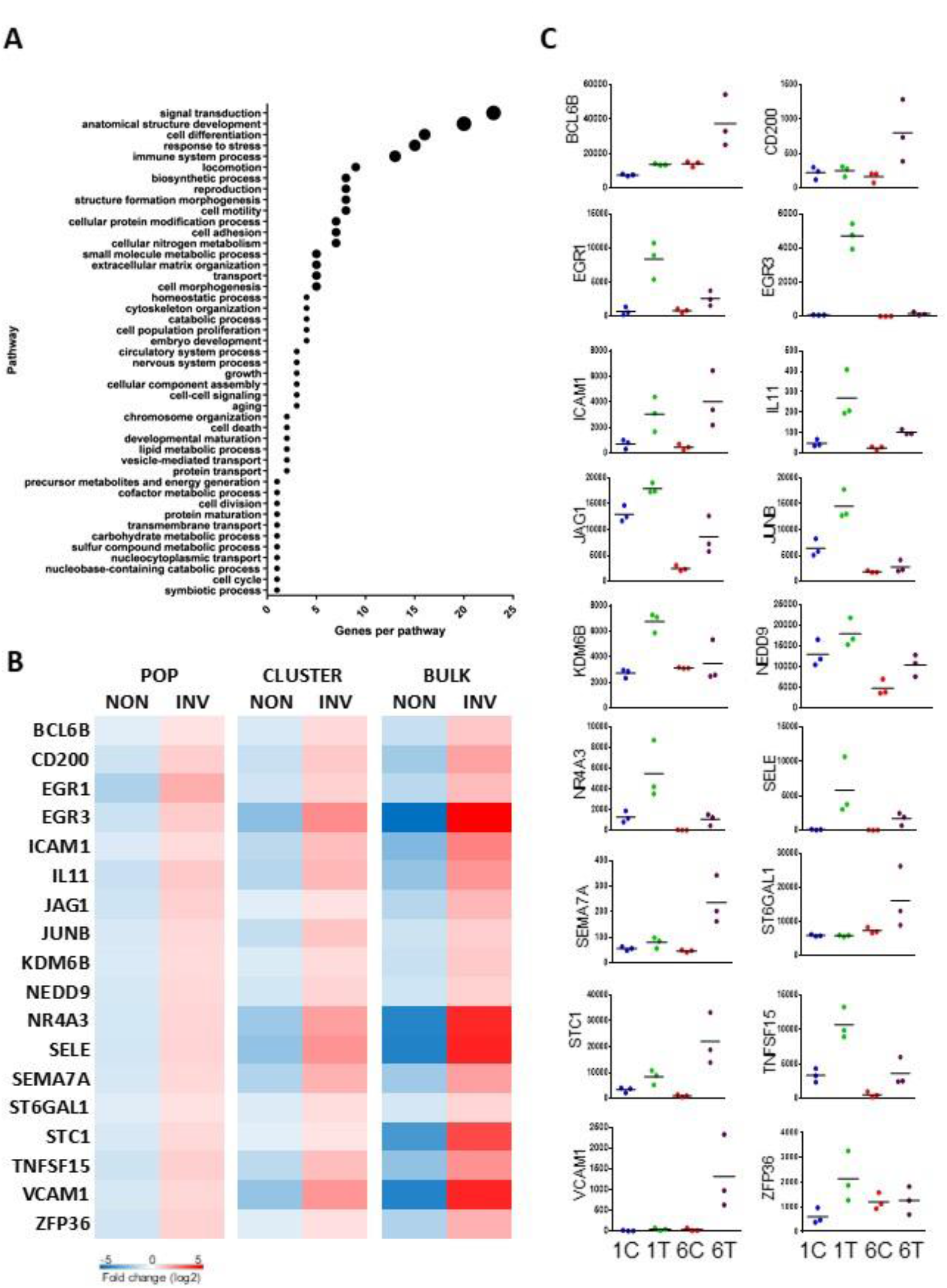
Analysis of gene expression patterns revealed the majority of candidate genes regulate signal transduction, developmental processes, and immune evasion. **(A)** GO Pathway analysis of 39 candidate genes upregulated with endothelial cell invasion. **(B)** Heatmap displaying expression of genes implicated in immune responses for Population, Cluster and Bulk RNA-seq Analyses. **(C)** RNA-seq data depicting upregulation of genes involved in immune responses occurs with endothelial activation. Samples were collected at 1 and 6 hr in the absence of activation (1C, 6C) and with activation (1T, 6T). All y-axes indicate relative mRNA expression levels, and data points show values observed from three independent donors. Black line indicates the average expression.

### Differentially Expressed Genes Generated from all Comparisons Showed High Congruency with Independent Experiments

We selected genes from each category listed in Table 1 to test for congruency of gene expression. Independent samples of INV and NON cells (6 hr incubation) were analyzed with qPCR (Fig. 4A). All gene candidates showed higher expression in INV compared to NON in at least one donor. No significant upregulation was observed for the CDH5 control. Bulk RNA expression is shown in Fig 4B to indicate independent expression in individual donors in control and treated groups at 1 and 6 hrs.

**Figure 4.**
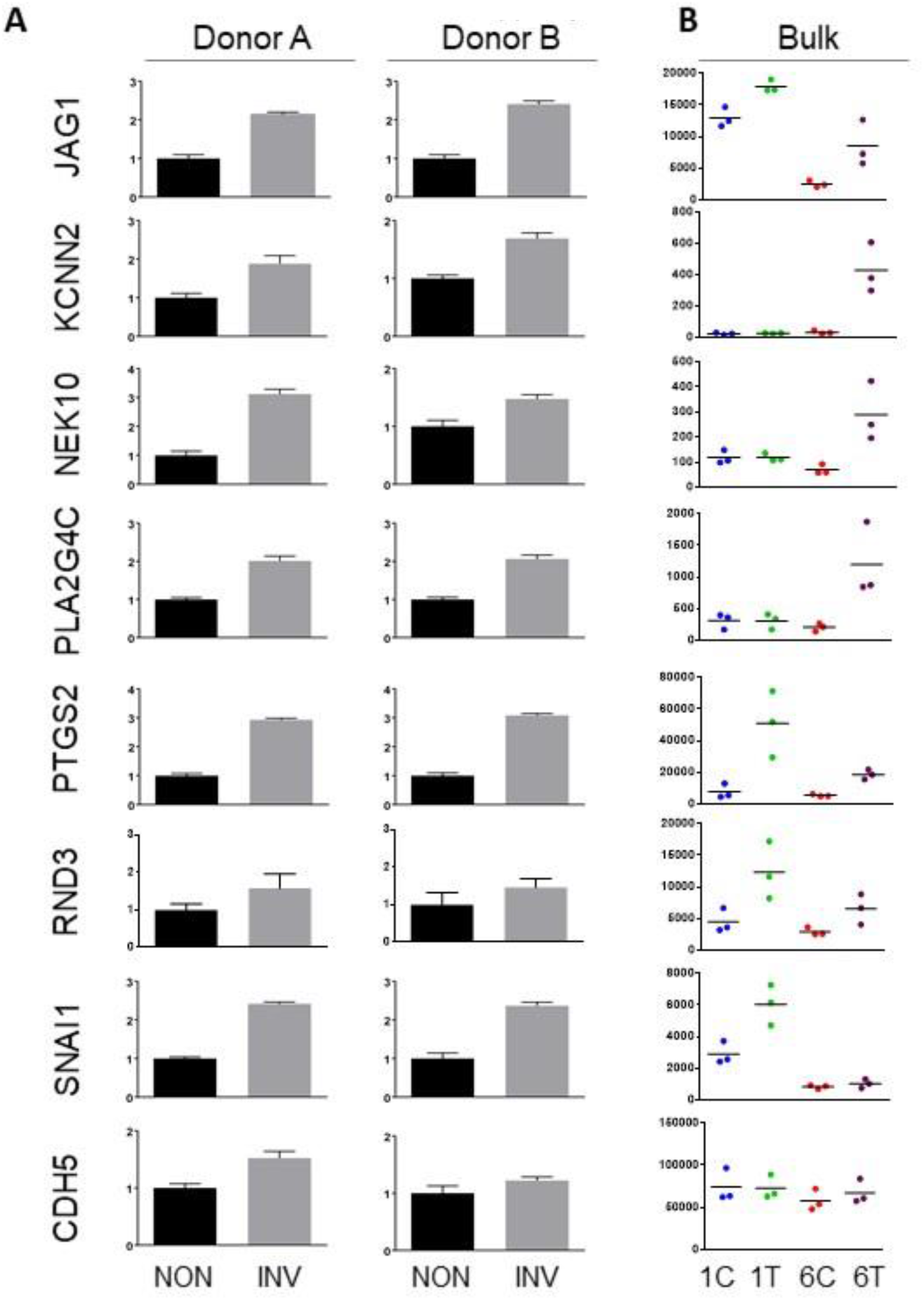
Validation of mRNA upregulation in genes from integrated analysis. Genes from each category in Table 1 were chosen for comparison. (**A**) qPCR for Donor A (left) and Donor B (right) from non-invading (NON) and invading (INV) samples. All y-axes indicate fold change represented as 2^-ΔΔct^. **(B)** Relative mRNA expression levels from bulk RNA-seq analyses. All y-axes indicate mRNA expression levels, and data points show values observed from three independent donors. Black line indicates average expression.

For further validation of transcriptomic data, we analyzed changes in protein expression of two upregulated genes that are known to be associated with angiogenic responses. Protein lysates collected from NON and INV ECs after 6 hrs of incubation were analyzed for PTGS2, SNAI1, and ERK1/2 (Fig. 5A). Normalization of PTGS2, SNAI1, and ERK1/2 to GAPDH or CD31 loading controls revealed significant upregulation of PTGS2 and SNAI1, but no change in ERK1/2 controls (Fig. 5B). These results were reinforced by immunofluorescence, showing increased staining intensity of PTGS2 and SNAI1 in invading cells compared to non-invading cells (Fig. 5C). Counterstaining with tubulin (red) allows visualization of sprouting structures. No increases in expression of the ERK1/2 control were seen in invading structures. These results confirmed that expression of PTGS2 and SNAI1 proteins increase with the onset of endothelial cell invasion.

**Figure 5.**
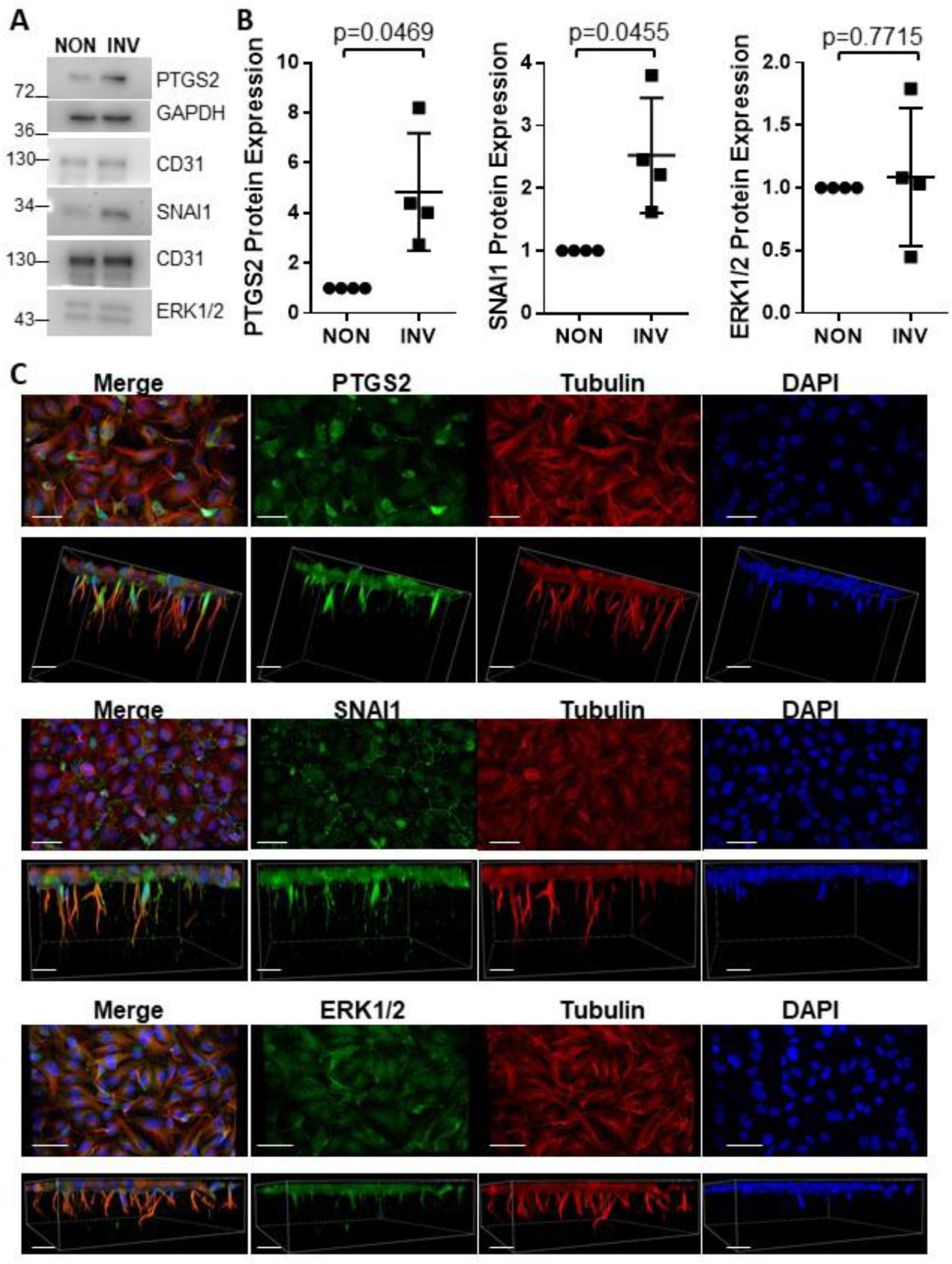
PTGS2 and SNAI1 protein expression increased in invading cells. Cells were allowed to invade for 6 hrs prior to analysis. **(A)** Western Blots comparing subpopulations of NON and INV cells were probed with antibodies directed to PTGS2, SNAI1, ERK1/2, and GAPDH or CD31 loading controls. **(B)** Quantification of band intensities normalized to controls from at least 4 independent experiments. Statistical values calculated with a Welch’s t-test. p=0.0469 (PTGS2),p=0.0455 (SNAI1), p=0.7715 (ERK1/2). (C) 3D projections of 6 hr invasion samples stained with PTGS2, SNAI1 and ERK1/2 antibodies (green), as well as tubulin (red) and DAPI (blue) to indicate invading structures and nuclei, respectively. Top panels show *en face* view of monolayer; bottom panels show side views of invading structures. Scale bars = 100 µm.

### SNAI1 Knockdown Reduced EC Invasion Distance

Because *SNAI1* has previously been reported to promote endothelial-mesenchymal transition (EndMT) during angiogenesis,^52,53^ we next tested for a role for *SNAI1* in initiating invasion. We compared siRNA-mediated knockdown of *SNAI1* (siSNAI1) to a beta 2 microglobulin (siβ2M) negative control. Previous studies using siβ2M reveal no negative effects on endothelial sprouting.^44,46,54–58^ Successful knockdown was observed for *SNAI1* (Figs. 6A and 6B) that was associated with modestly reduced invasion density (Fig. 6C) and significantly reduced invasion distance (Fig. 6D), consistent with a prior report.^59^ *SNAI1* expression is upregulated rapidly at the onset of sprout initiation in invading cells (Fig. 6B) and is required for optimal invasion responses, providing a correlation between rapid induction of SNAI1 and angiogenesis.

**Fig. 6.**
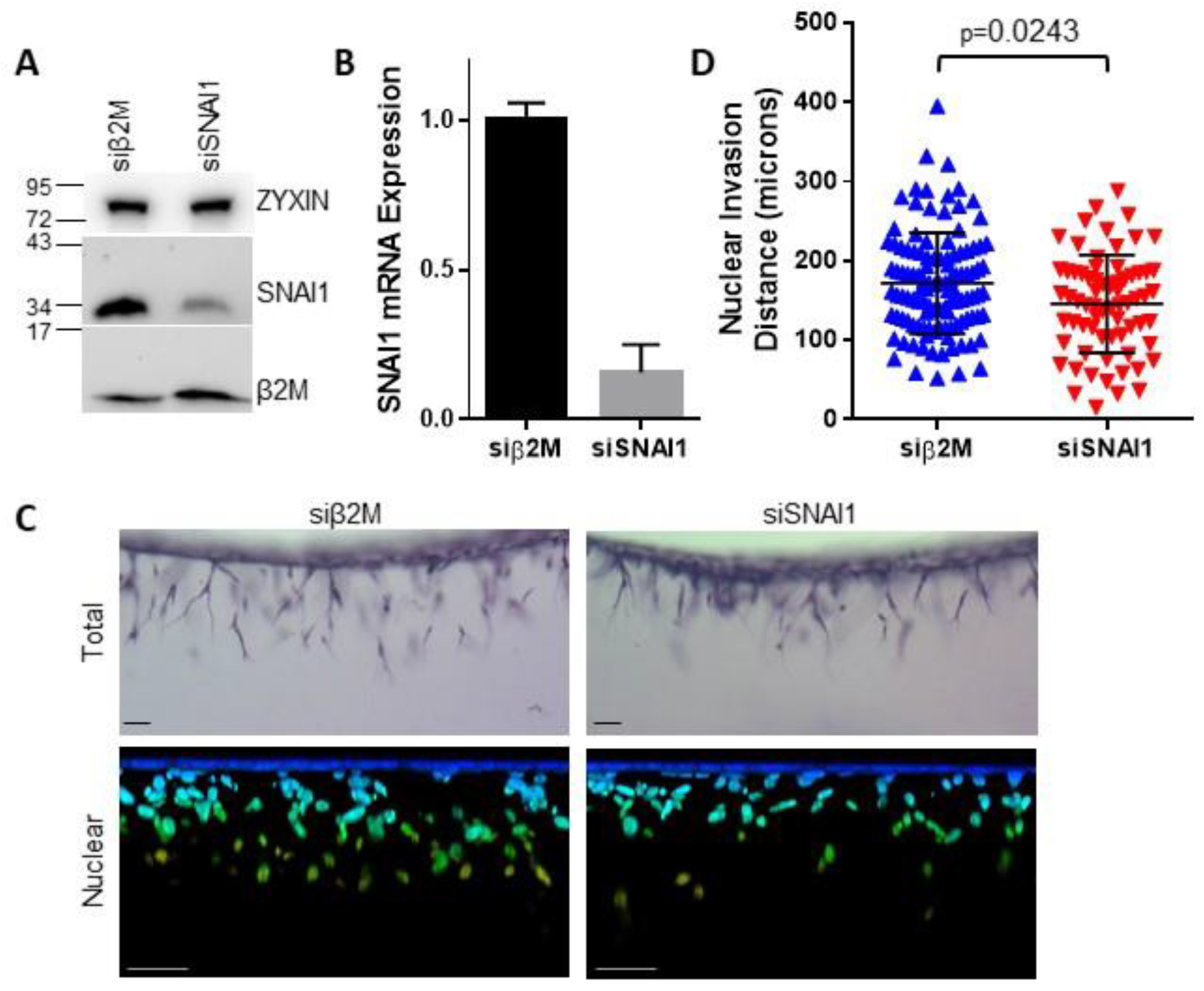
SNAI1 knockdown reduced endothelial cell invasion distance. Cells were treated with siRNA to knock down β2M (siβ2M control) and SNAI1 (siSNA1) and allowed to invade. Gene silencing was confirmed by **(A)** Western blotting of protein lysates (6 hr) probed with antibodies indicated. ZYXIN serves as a loading control. **(B)** qPCR analysis of RNA templates in siRNA treated cells. **(C)** Cells were allowed to invade into collagen matrices for 24 hrs. Upper panels: Side view images of siβ2M control and siSNAI1 samples stained with toluidine blue to show development of the sprouting structures. Lower panels: 3D renderings of DAPI stained samples captured with confocal microscopy. Colors represent Alpha Depth Coding to visualize depth of nuclear invasion. Blue, original monolayer; orange, deepest invasion. Scale bars = 100 µm. **(D)** Quantification of nuclear invasion distance. Representative experiment shown (n=3); At least 50 structures were analyzed from three experiments. p=0.0243, using Mann-Whitney U test.

### Loss of JunB Reduced EC Invasion Density and Distance

JUNB is an AP-1 transcription factor that has recently been implicated in retinal angiogenesis.^60^ To determine if changes in JUNB protein expression follow transcriptional upregulation, endothelial cells invading into collagen matrices for 3 hrs were stained with antibodies to JUNB and tubulin. After categorizing cells as invading or non-invading based on tubulin staining (Fig. 7A), JUNB nuclear intensity was normalized to DAPI intensity. The results showed that invading endothelial cells have significantly higher expression of JUNB compared to non-invading cells (Fig. 7B). Because silencing of *JUNB* was inefficient when using siRNAs (data not shown), antisense oligonucleotides (ASOs) were designed to target *JUNB*. Targeting *JUNB* significantly reduced JUNB protein (Fig. 7C) and mRNA expression (Fig. 7D). As expected, loss of *JUNB* resulted in a significant decrease in invasion density (Figs. 7E and 7F) and invasion distance (Figs. 7E and 7G). These data confirm *JUNB* is required for successful invasion responses and agree with a prior report.^60^

**Fig. 7.**
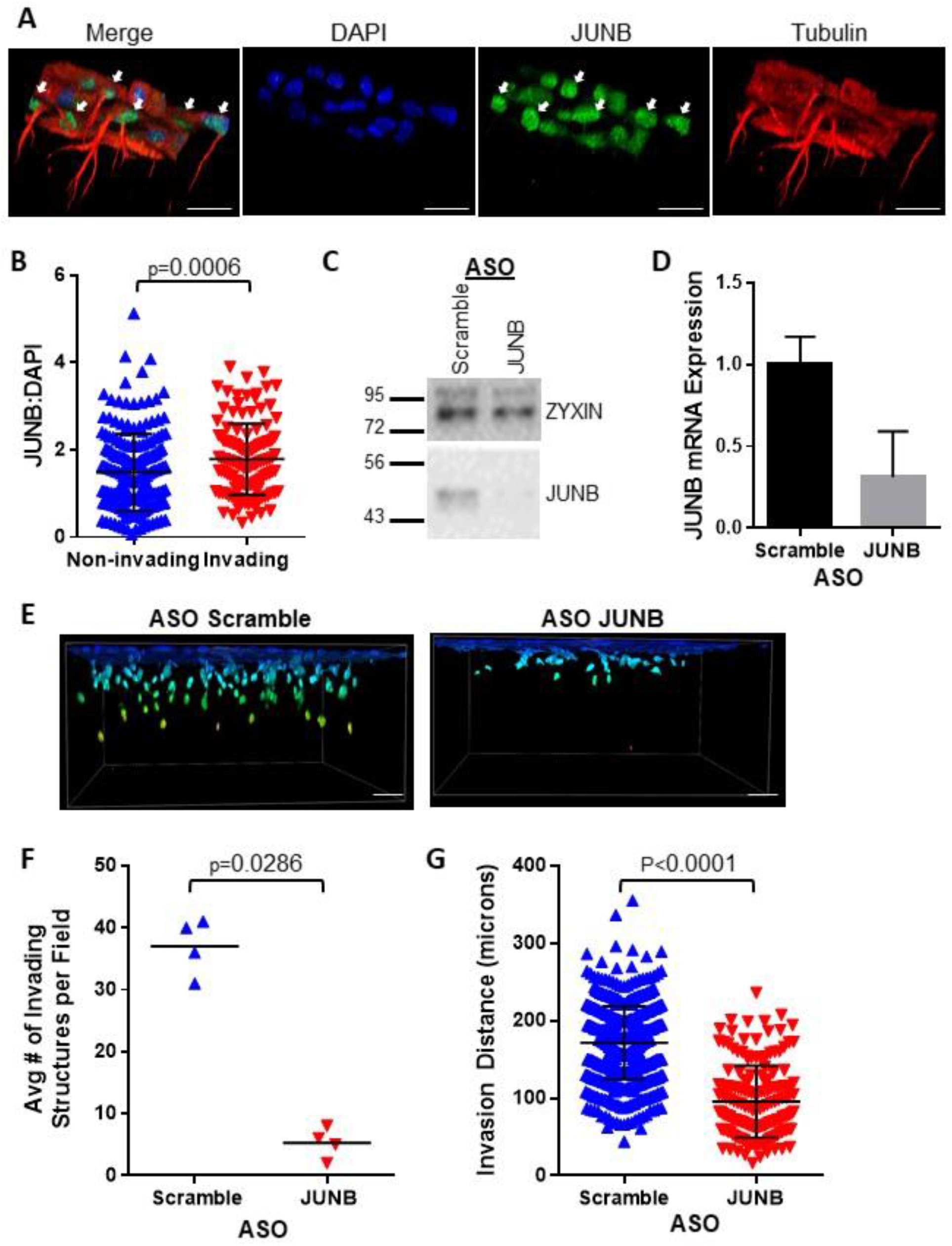
JunB is required for EC invasion responses. **(A)** JUNB nuclear intensity was determined using immunofluorescence. Samples were fixed at 3 hrs and stained for tubulin (red), JUNB (green), and DAPI (blue). **(B)** Quantification was performed as described in Materials and Methods in at least two independent experiments. Data shown are average JUNB nuclear intensity normalized to DAPI. White arrows depict invading cells. p=0.0006 with Mann-Whitney U test. **(C-G)** Silencing of JUNB using antisense oligonucleotides (ASO). **(C)** Representative western blot (n=4) determining JUNB expression using ASO specific to JUNB or scrambled control. Changes in protein expression were confirmed using antibodies directed to JUNB or ZYXIN as a loading control. **(D)** qPCR analysis showing relative JUNB expression. **(E)** Cells were allowed to invade into collagen matrices for 24 hrs. Side view of 3D images of scrambled and JUNB ASO. Samples were stained with DAPI and analyzed with confocal microscopy. Alpha Depth Coding was used to visualize depth of nuclear invasion. Scale bars = 100 µm. **(F)** Quantification of invasion responses. Representative experiment from 3 independent experiments. Data shown are average numbers of invading cells per field. Statistical significance was determined using Mann-Whitney U test; p=0.0286. **(G)** Quantification of nuclear invasion distance. Representative experiment shown (n=3); At least 75 structures were analyzed from three independent experiments. P<0.0001, using unpaired Student’s t-test.

### RND3 Contributed to EC Sprout Initiation

Sprout initiation is associated with changes in endothelial cell motility, and multiple genes involved in regulating these processes were upregulated. Although *RND1* and *SEMA7A* have previously been shown to play a role in endothelial motility and angiogenesis,^61,62^ few data are available on the role of *NEDD9*, *NUAK2*, *RAPH1*, or *RND3*. RND3 is an atypical Rho GTPase (also known as RhoE) that has previously been implicated in cell rounding, loss of stress fibers^63,64^ and barrier function.^65,66^ Thus, we performed siRNA-mediated silencing to determine if *RND3* was required for endothelial cell invasion. siRNA directed to *RND3* significantly reduced mRNA expression compared to control treatment (Fig. 8A). *RND3* silencing also resulted in a noticeable reduction in nuclear invasion distance (Fig. 8B and 8C) but no change in invasion density (Fig. 8D). A closer analysis of the morphology of invading cells revealed shorter overall processes and filopodia that appeared to be stunted (Fig. 8E). Quantitation of filopodia length revealed a significant decrease with siRND3 treatment compared to siNEG control (Fig. 8F). These data agree with the original report of RND3 overexpression driving filopodia formation^67^ and suggest RND3 upregulation may be necessary for filopodia extension, which is known to be required for tip cell migration^68^.

**Fig. 8.**
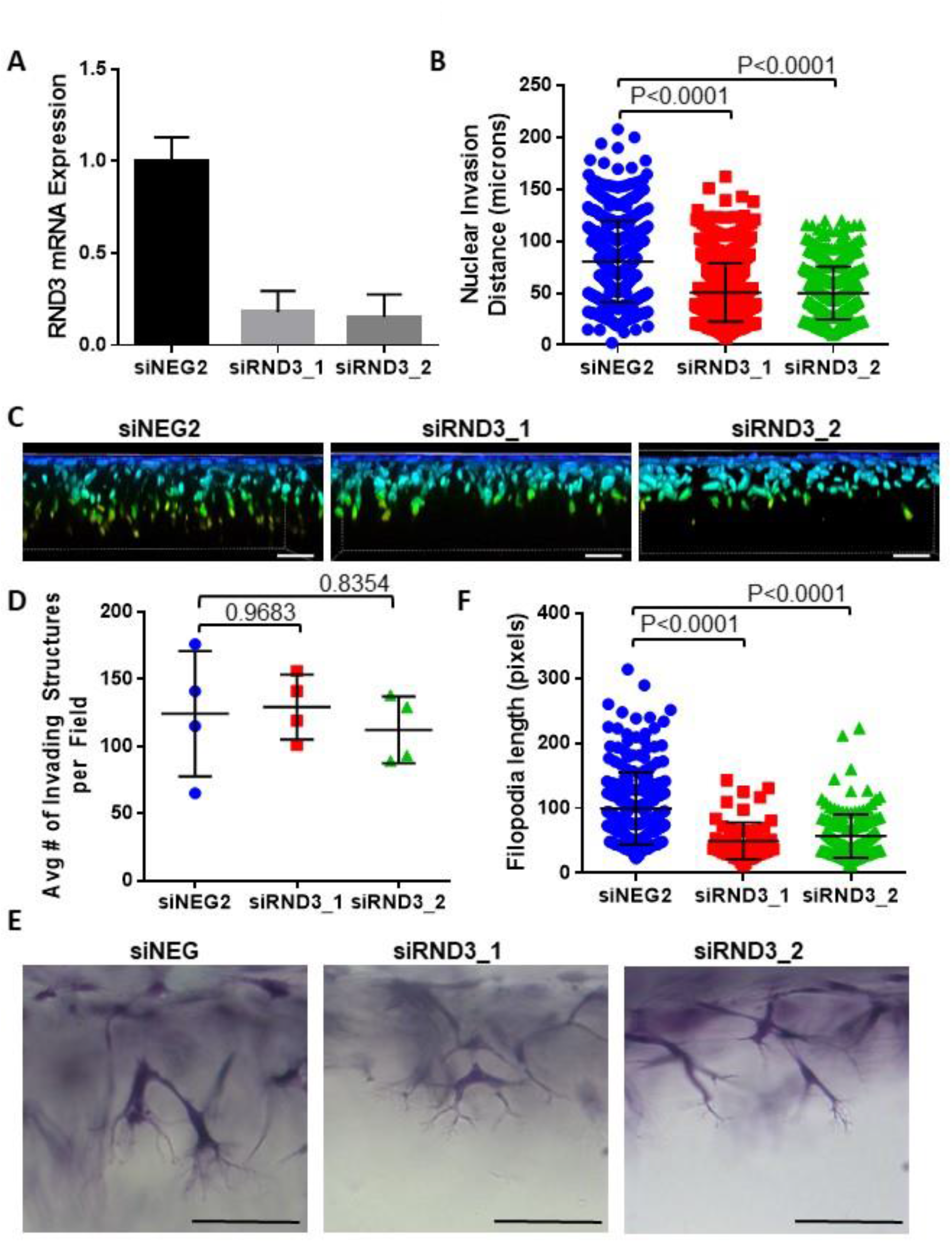
RND3 fine tuned filopodia to control EC invasion responses. Silencing of RND3 using siRNA was performed prior to placing cells in 3D invasion assays. **(A)** qPCR analysis showing relative RND3 expression in negative control (siNEG2) and two separate siRNAs targeting RND3 (siRND3_1 and siRND3_2). Data shown are from a representative experiment (n=3). **(B)** Quantification of nuclear invasion distance. At least 400 structures were analyzed from 3 experiments. p=0.0001, using one-way ANOVA. Representative experiment shown. **(C)** Samples (24 hr) were stained with DAPI and analyzed with confocal microscopy. Alpha Depth Coding was used to visualize depth of nuclear invasion. Scale bar = 100 µm. **(D)** Quantification of invasion responses (n=3). Data shown are average numbers of invading cells. Statistical significance was determined using Kruskal-Wallis p=0.6024. **(E)** High magnification images of invasion responses observed with siNEG2 and siRN3D_1 and siRND3_2. Samples were stained with toluidine blue. Scale bar = 100 µm. **(F)** Quantification of filopodia length. Calibrated images were used to measure filopodial lengths from at least 25 invading structures in 3 experiments. P<0.0001, using one-way ANOVA test.

To confirm a role for RND3 in filopodial extension *in vivo*, we used mice heterozygous for *Rnd3*. Haplosufficient *Rnd3* mice were necessary because homozygous mice are born at a frequency of 0.9% of live births.^47^ No significant reductions were seen in retinal outgrowth between wild type (WT) and *Rnd3^+/-^* P5 mice (Figs. 9A and 9B). To analyze filopodial extensions, additional analyses focused on the retinal front at P5, where it appeared filopodial extensions were shorter in *Rnd3^+/-^* tip cells (Fig. 9C). Quantification of filopodial length revealed that *RND3^+/-^* mice displayed significantly shorter filopodia than WT littermates (Fig. 9D), confirming reduction in filopodia length observed *in vitro* (Fig. 8). Perhaps not surprisingly, since *Rnd3^+/-^*mice retain one copy of *Rnd3*, this may explain why no differences in retinal outgrowth are observed. Regardless, *Rnd3* upregulation appears to be required for filopodial extension, which is a key component of successful angiogenic responses.^68^

**Fig. 9.**
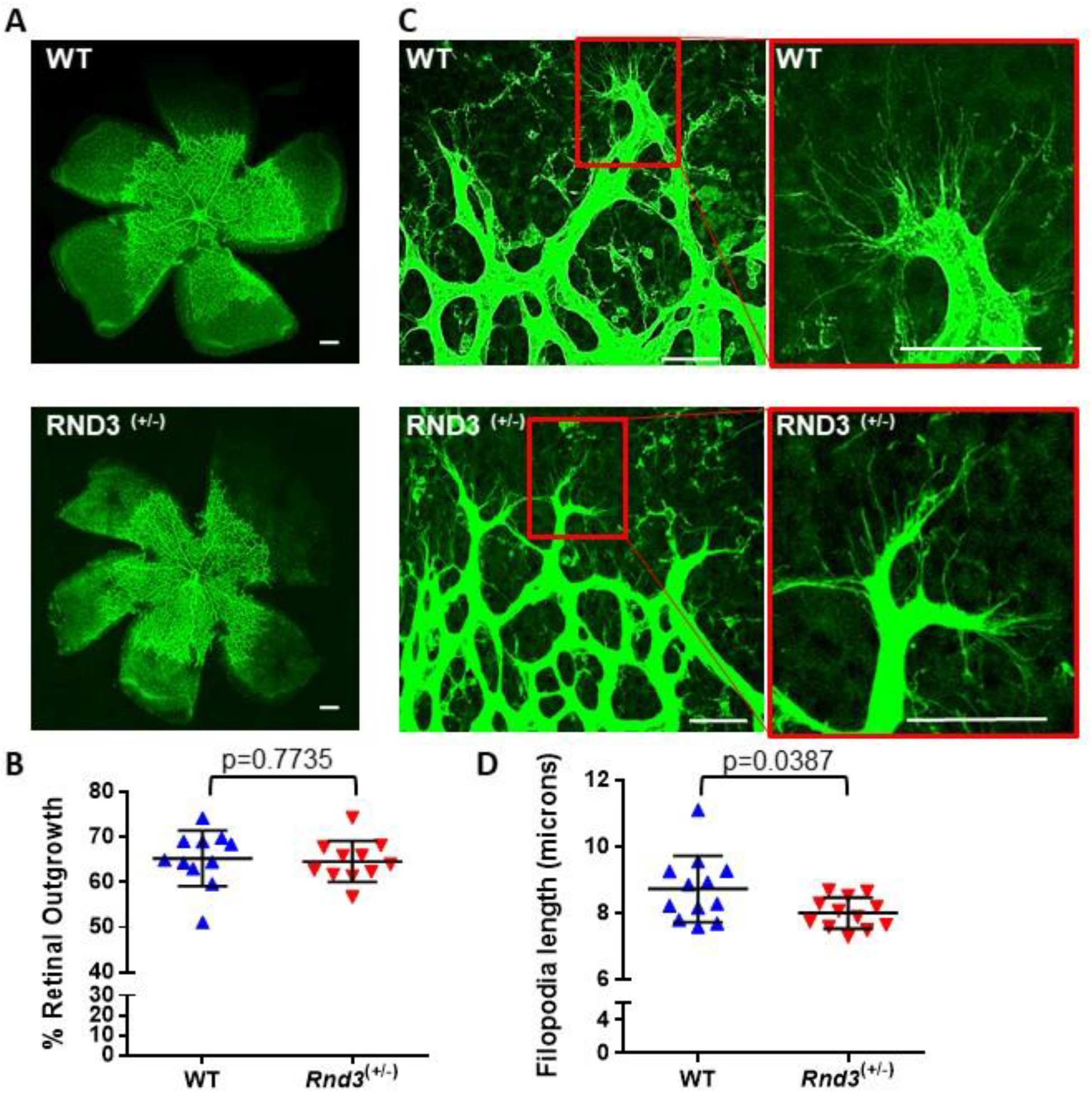
RND3 haploinsufficiency mimics resulted in shorter tip cell filopodia. Retinas from postnatal day 5 (P5) were collected from wild type (WT) and *Rnd3*^+/-^ littermates (n=6 each) and stained with fluorescent isolectin B4. (**A**) Photographs of P5 retinas. Scale bar = 250μm. **(B)** Quantification of percent retinal outgrowth at P5. Data from individual eyes in each group are shown. Percent outgrowth calculated as edge of vascular front normalized to lobe length, p=0.7735 using Mann-Whitney U test **(C)** Images of the retinal angiogenic front at P5. Magnified views (red boxes) illustrate tip cell filopodia seen in WT and *Rnd3*^+/-^littermates at P5. Scale bars = 20 µm. **(D)** Quantification of filopodia length in tip cells at P5. Data represent average filopodia length from each retina. p=0.0387, using Mann-Whitney U test. Data were collected from 4-6 images per retina; images contained an average of 12 (or between 5-22) tip cells.

## DISCUSSION

This study analyzed multiple independent biological replicates with three transcriptomic analyses to reveal gene expression changes that distinguish invading from non-invading endothelial cells, and it provides a new set of markers to characterize mRNAs that are upregulated at the onset of angiogenic sprouting. Using combined scRNA-seq Population and Cluster Analyses, along with bulk RNA-seq, we discovered 39 candidate genes in common that were rapidly upregulated within 6 hrs of initiating invasion. The majority are associated with morphogenesis or developmental processes, supporting a pronounced and rapid change in phenotype at the onset of angiogenic sprout initiation. Our findings suggest approximately one-half of the upregulated genes regulate immune responses or are associated with immune evasion, providing evidence that significant phenotypic changes occur in endothelial cells within hours after activation and sprout initiation. We validated a role for three of these genes (*SNAI1, JUNB, RND3*) in our 3D model by demonstrating that silencing resulted in significantly impaired cell invasion. Notably, *RND3* knockdown reduced filopodia length. Additionally, we successfully verified with qPCR, congruency of expression of a subset of our list of 39 in independent invading and non-invading samples. Furthermore, we validated an upregulation of protein expression in invading cells for SNAI1, PTGS2 and JUNB. Finally, we show in a mouse retinal model that haploinsufficiency of *Rnd3* retarded filopodia growth. These data provide a new set of markers to characterize early sprouting events in angiogenesis.

Our data reveal a role for RND3 as an early marker of angiogenic sprouting that enhances filopodia extension and endothelial cell migration in 3D. We found that silencing RND3 expression shortened filopodia length and reduced invasion distance *in vitro*. Shortened filopodia are consistent with early reports that indicate RND3 overexpression promoted substantial filopodial extensions.^69^ A connection between filopodia, which are prominently displayed on tip cells and cell migration has been established. The absence of filopodia slowed tip cell migration during retinal angiogenesis.^70^ More recently, endothelial cell filopodia were demonstrated to rapidly and actively perceive the environment and accelerate decisions for tip cell differentiation,^2,71,72^ and filopodia are needed for faster searching,^73,74^ as loss of filopodia led to a slower selection of tip cells.^68,72,75–77^ Together, these reports link filopodia extensions with tip cell motility. Although we did not observe a decrease in the speed of the outgrowth of retinal angiogenic vessels in P5 mice with *Rnd3* haploinsufficiency, this may be due to incomplete loss of endothelial *Rnd3* expression. A role for *Rnd3* is supported by decreased density of cardiac vessels in *Rnd3^+/-^* mice compared to wild type littermates with challenge using pressure overload,^78^ so perhaps challenge is necessary. Additional studies are needed, ideally with endothelial-specific silencing of *Rnd3* to prove its requirement in retinal angiogenesis *in vivo*. RND3 has been linked to invasive behavior in cancer cells^79^ fitting with the data from our 3D model of endothelial invasion, but contrastingly independent of a filopodial phenotype. However, RND3 can also impair stress fiber formation and induce cell cycle arrest.^64^ It remains to be determined if RND3 regulates additional steps in angiogenesis beyond filopodial extension.

A substantial number of upregulated transcripts (18 of 39 genes) are associated with immune responses and immune evasion. Angiogenesis is well-accepted to occur alongside inflammation.^80^ Resident cells such as macrophages, dendritic cells, lymphocytes, and microglia promote angiogenesis and anastomosis during zebrafish development,^81^ collateral vessel growth,^82^ macular degeneration,^83^ chick CAM assays,^84,85^ retinal angiogenesis,^81^ and developmental and pathological angiogenesis.^86,87^ Leukocytes secrete angiogenic factors, including chemokines and VEGF. The presence of these pro-angiogenic factors promotes endothelial growth and migration toward the initial source (i.e. leukocytes), and direct interactions have been documented elegantly by Ruhrberg and colleagues.^81^ Others have shown angiogenic endothelial cells navigate the extravascular environment often encountering macrophages, monocytes, CD45+ cells, and dendritic cells.^81,88–90^. It makes sense that endothelial cells should be equipped to display the proper cell surface markers to avoid immune recognition by individual resident immune cells. In addition to well-characterized markers of endothelial activation such as SELE, ICAM1, VCAM1, two prominent candidates are identified here: CD200 and stanniocalcin 1 (STC1). STC1 has long been known to correlate with angiogenic responses, but its exact function has remained elusive.^91^ Recently, STC1 was shown to promote intracellular retention of calreticulin (CALR), an “eat me” signal, and promote immune evasion in liver cancer.^92^ It is worth noting that both *STC1* and *CALR* are highly abundant endothelial transcripts (see Supplemental Table 6). In addition to STC1, CD200 is well-described in the cancer literature to mediate immune evasion.^93^ CD200 and multiple receptors are also expressed by invading trophoblasts and tissues in the decidua in mouse and humans,^94^ where they are anticipated to promote immune tolerance in pregnancy.^95^ Thus, our data indicate that rapid upregulation of *STC1* and *CD200* expression along with well characterized markers of endothelial activation occur to allow ECs to avoid immune recognition and activation of resident immune cells as angiogenesis proceeds.

Our results show that collagenase treatment to dissociate cells artifactually activates *FOS* expression in endothelial cells. These findings reinforce recent reports that collagenase dissociation upregulates the expression of early response genes and affects clustering by exaggerating differential gene expression^50,51,96^. In the prior reports, digests were typically performed for one hour, whereas in this study, cells were exposed to collagenase for 10 minutes. Regardless, we collected cells treated with and without collagenase and observed a 3-fold increase in FOS expression only in collagenase-treated cells compared to non-treated cells. It is possible that we observed a minimal impact of collagenase digestion on gene expression because of the limited exposure time to collagenase compared to longer digestion times in other studies.^50,51^ Based on the observed FOS upregulation with collagenase treatment (Supplemental Fig. S3A), cells with high FOS expression were excluded to avoid skewing Cluster Analysis. In addition, we confirmed no significant upregulation of candidate genes occurred in response to collagenase treatment (Supplemental Fig. S3B). Our studies reveal that even limited treatment with collagenase can upregulate the FOS early response gene and potentially skew clustering results.

The Clustering Analysis aided an unbiased analysis of sprouting responses in 3D. Clustering Analysis is inherently dependent on input parameters and can be highly sensitive to threshold and filtering.^97^ When separating the total cell sample into two clusters, the results aligned with the independent results from the Population Comparison and bulk RNA-seq analyses. The results of the Clustering Analysis are further strengthened by the reproducibility within the non-invading and invading cluster profiles of both biological replicates. In agreement with an early phenotypic change, several tip cell markers were significantly upregulated in the Cluster Analyses. Angiogenic tips cells are characterized as highly migratory cells that extend multiple filopodia and lead the way during new blood vessel growth.^98^ The integrated comparison that evaluated invading and non-invading endothelial cells (Table 1) revealed significantly higher expression of known tip cell markers ADAMTS1,^99^ JUNB,^60^ BCLCB, RND1, VCAM1^100^ in invading endothelial cells. The abundant literature reporting tip cell markers has been collected using a variety of models.^32,101–104^ Often the new blood vessel growth studied in these models is at an advanced stage of progression, where tip, stalk, and phalanx cells are present.^4,27,31,32^ In the current study, endothelial cells are still in the early or nascent stages of emerging from an existing monolayer. Although cells extend filopodia into the collagen matrix, junctions are intact, and nuclei largely remain on the surface of the collagen matrix, raising the possibility that all cells in the invading population are not yet fully differentiated or perhaps committed, because endothelial cells are highly plastic and oscillate between tip, stalk, and other cell phenotypes.^31,75,77,105^ These data indicate the existence of an emerging tip cell population in the 3D model of endothelial invasion, based on upregulation of multiple tip cell markers in both the Population and Cluster Analyses. Further study is needed to address a functional role for the previously unidentified candidate genes in controlling tip cell behaviors.

Our final list of 39 candidate genes is consistent with EndMT occurring at the onset of invasion. The transitory state of partial EndMT has been proposed as a method for endothelial cells to acquire invasive properties while still maintaining an endothelial identity.^53^ SNAI1, a known activator of EndMT, had a significant impact on endothelial invasion distance, as knockout cells exhibited impaired migration when compared to controls. The Hughes laboratory has shown that SNAI1 and Slug expression increase with time during endothelial cell invasion in 3D fibrin matrices and are essential for endothelial cell migration and sprouting, concluding that they do not act redundantly.^59^ We observe low levels of Slug expression, perhaps because of differences in experimental conditions (the addition of serum or phorbol ester), the 3D matrix used (fibrin vs. collagen), or timing (6 hrs of invasion vs. 3-6 days). The data shown here indicate that SNAI1 is activated quickly (1 hr) on collagen matrices and that exposure to S1P and growth factors amplifies this upregulation. Other genes upregulated with invasion that have been previously implicated in EndMT include *NDST1*,^106^ *RND3*,^107^ *SEMA7A*,^108^ *STC1*,^109^ and *VCAM1*^110^ reinforcing an EndMT phenotype at the onset of angiogenic sprouting.

Our data raise the possibility that AP-1 transcription factors promote a migratory phenotype consistent with EndMT. We observed upregulation of the AP-1 transcription factor *JUNB* in invading cells. AP-1 transcription factors have been shown to be required for vascular development in the embryo, placenta, and retina. ^60,111–113^ Specific evidence for a requirement in endothelial cells has been provided by both constitutive^114^ and inducible^60^ endothelial knockouts. These findings agree with the previously reported opening of AP-1 transcription sites and a key role for *JunB* in retinal angiogenic endothelial cells that dive into the deep retinal layer^60^. Consistent with that report, we knocked down *JUNB* using ASOs and found impaired invasion, quantified by reduced invasion distance and density of activated endothelial cells. Given that an AP-1 binding site exists in the promoter region of *SNAI1*^115^ and *Slug*,^116^ AP-1 transcription factors may be rapidly upregulated to direct the onset of EndMT.

Interpreting data from large transcriptomic experiments can be daunting, as endless lists of differentially expressed genes are often generated. Consistent with others,^31^ we employed a three-tiered analysis approach using scRNA-seq and bulk RNA-seq data from five individual donors in which invading cells were efficiently separated from non-invading cells. From these analyses, we identified 39 candidate genes upregulated in activated, invading endothelial cells. Evaluation of these genes highlights a stark and rapid change in plasticity that occurs in endothelial cells as diverse transcriptional pathways are activated to accomplish rapid, coordinated convergence leading to the initiation of angiogenic sprouting.

## Acknowledgments

Research reported in this publication was supported by the Texas A&M University T3 grant 02-247008 (KJB) and the Department of Molecular and Cellular Medicine. We thank Drs. Peng Yu and Zhengyu Guo of the TEES-AgriLife Center for Bioinformatics and Genomic Systems Engineering for assistance with bulk RNA-seq analysis and annotation.

## DISCLOSURES

### Authors’ contributions

Experimental design: KJB, CAA, CLD, and JC

Data collection: CAA, ZC, YC, SR, KJB, CLD, and AC Data Analysis: CAA, AC, and KJB

Manuscript drafting: KJB, CAA, GBW, JC, SC, and AC

Manuscript editing and approval: all authors

## AVAILABILITY OF DATA AND MATERIAL

All data files used for analyses are included in supplemental materials.

### Code availability

Single-cell RNA sequencing analyses using Cell Ranger were performed by Bioinformatics Core at Texas A&M Institute for Genome Sciences & Society (TIGSS). Projections were generated using 10X Genomics Loupe 6.0.1

## Supplemental Figures

**Supplemental Fig. S1 Uncropped western blots used in preparation of the manuscript. (A)** Blots from Figure 5A. **(B)** Blots from Figure 6A (SNAI1 exp). **(C**) Blots from Figure 7C. (JUNB ASO)

**Supplemental Fig. S2 Photographs of samples collected when non-invading and invading endothelial populations were separated from three-dimensional assays for single-cell sequencing. (A)** Photographs of *en face* views showing cell monolayers on the surface of collagen matrices before and after trypsinization. **(B)** High magnification photographs of a side view of invading samples before and after trypsinization. **(C)** Photographs of non-invading (NON) and invading (INV) cells liberated by trypsin and collagenase treatments, respectively, submitted for scRNA-seq analyses.

**Supplemental Fig. S3 Collagenase disassociation artificially induced FOS expression but not our candidate genes**. At 6 hr of invasion, collagen matrices and associated cells were directly lysed in RLT buffer or digested with collagenase and then lysed in RLT buffer before RNA extraction. Y-axis represents fold change in expression between non collagenase treatment and collagenase treatment for the (**A**) early response genes shown and (**B**) a subset of candidate genes.

**Supplemental Table S1. Primer sequences used in qPCR.**

**Supplemental Table S2. scRNA-seq metrics.** Organized by sample for each cell lot and cell population, data shows estimated number of cells sequenced, fraction of reads in cells, mean reads per cell, median genes per cell, total genes detected and the median UMI counts per cell.

**Supplemental Table S3. Differentially expressed genes from Population Comparison between non-invading and invading cells for aggregated donors A and B.** Differentially expressed genes are indicated with Ensembl ID, gene name, average expression of non-invading, average expression of invading, log 2 fold change, and p-value.

**Supplemental Table S4. Differentially expressed genes from Cluster Analysis for aggregated donors A and B.** Differentially expressed genes are indicated with Ensembl ID, gene name, average expression of non-invading, average expression of invading, log 2 fold change, and p-value.

**Supplemental Table S5. Bulk RNA-seq metrics.** Organized by sample for each cell lot and time point, data shows the number and percent of uniquely mapped reads.

**Supplemental Table S6. Genes significantly upregulated from bulk RNA-seq analysis between 1 HR CON, 1 HR TX, 6 HR CON, and 6 HR TX samples.** Organized alphabetically by hit ID and location to reference genome hg19. Base mean for the 3 replicates with A belonging to the first sample in the comparison column heading and B to the second, followed by fold change, log 2 fold change, p value (pval), padj value and comparison.

**Supplemental Table S7. Overlap of genes significantly upregulated in the Population Comparison, Cluster Analysis, and Bulk RNA-seq in relation to each other.** Table showing genes commonly upregulated in all 3 transcriptomic data sets. The first two column shows the upregulated genes in the Population Comparison and Cluster Analysis, respectively. The 3^rd^ shows the list common to Population and Cluster Analyses. The bulk RNA-seq hits are listed in the next column. The last column indicates the overlap within the Population Comparison, Clustering Analysis, and bulk-RNA seq.

**Supplemental Table S8. Known roles of candidate genes consistently upregulated in invading cells.** Genes are listed alphabetically in functional categories based on GO terms, identified by gene symbol and name. Genes in bolded text have not been shown to play a role in angiogenic responses.

